# Crif1 Promotes Osteoporosis in Mice after Radiation

**DOI:** 10.1101/725408

**Authors:** Lixin Xiang, Li Chen, Yang Xiang, Fengjie Li, Xiaomei Zhang, Yanni Xiao, Lingyun Zou, Jiang F. Zhong, Shengwen Calvin Li, Qian Ran, Zhongjun Li

## Abstract

Radiation induces rapid bone loss and enhances bone resorption and RANKL expression. RANKL provides the crucial signal to induce osteoclast differentiation and plays an important role in bone resorption. However, the mechanisms of radiation-induced osteoporosis are not fully understood. Here, we show that Crif1 expression increases in bone marrow cells after radiation. Conditional Crif1 deletion in bone marrow cells causes decreases in RANKL expression and the RANKL/OPG ratio, and relieves bone loss after radiation in mice. We further demonstrated *in vitro* that Crif1 promotes RANKL secretion via the cAMP/PKA pathway. Moreover, protein-protein docking screening identified five compounds as Crif1 inhibitors; these compounds dramatically suppressed RANKL secretion and CREB phosphorylation when cells were exposed to forskolin. This study enriches current knowledge of the pathogenesis of osteoporosis and provides insights into potential therapeutic strategies for osteoporosis treatment.

## Introduction

Osteoporosis is a high-incidence disease closely associated with radiotherapy, hormonal status, age and glucocorticoid treatment. It is characterized by an imbalance in skeletal turnover with reduced bone formation and enhanced bone resorption, leading to an increased risk of bone fracture(Khosla & Hofbauer, 2017; Phetfong et al., 2016). Osteoblasts are the bone-forming cells derived from bone marrow mesenchymal stem/stromal cells (BM-MSCs) and play an important role in the regulation of bone mass. Meanwhile, osteoclasts are the principal cells capable of resorbing bone and play an essential role in bone remodelling(Rachner, Khosla, & Hofbauer, 2011). Exposure to radiation, such as radiotherapy for cancer, can cause rapid mineral loss and increase the number of osteoclasts within metabolically active, cancellous bone tissue, leading to structural deficits. However, the mechanisms of radiation-induced bone loss are not fully understood. Current treatment of osteoporosis is based mainly on inhibiting bone resorption or stimulating bone generation to increase bone mass, novel treatment strategies that have a dual mechanisms should be developed.

Osteoclasts are large, multinucleated cells derived from haematopoietic progenitors of the monocyte macrophage lineage(Boyle, Simonet, & Lacey, 2003). Their differentiation is mainly regulated by macrophage colony-stimulating factor (M-CSF), RANK ligand (RANKL), and osteoprotegerin (OPG)(Teitelbaum, 2000). M-CSF is required for osteoclast survival and proliferation, but RANKL and OPG play central roles in the activation of osteoclastogenesis(Boyce & Xing, 2007). By binding to its receptor RANK (on haematopoietic progenitors), RANKL provides the crucial signal to induce osteoclast differentiation from haematopoietic progenitor cells as well as to activate mature osteoclasts. OPG is a soluble decoy receptor that can bind to RANKL and negatively regulate RANKL binding to RANK(Wada, Nakashima, Hiroshi, & Penninger, 2006). BM-MSCs, osteocytes, osteoblasts, adipocytes, and activated T and B lymphocytes are the main sources of RANKL secretion. RANKL expression is promoted by radiation, inflammation, cytokines, hormones and a number of other agents, including those that signal through cyclic adenosine monophosphate (cAMP)/protein kinase A (PKA), glycoprotein 130 (gp130) and vitamin D receptor (VDR)(Martin & Sims, 2015; Nakashima et al., 2011).

Crif1 is a multifunctional protein that can interact with many proteins to induce cell cycle arrest, modulate oxidative stress, and regulate transcriptional activity through interactions with the DNA-binding domains of transcription factors(Chen et al., 2017; Chung et al., 2003; Kang, Hong, Kim, & Bae, 2010; Park et al., 2005; Ran et al., 2014). It is also the constitutive protein of the large mitoribosomal subunit required for the synthesis and insertion of mitochondrial-encoded OxPhos polypeptides into the mitochondrial membrane(Greber et al., 2015). Crif1 deficiency in macrophages impairs mitochondrial oxidative function and causes systemic insulin resistance and adipose tissue inflammation(Jung et al., 2018). Our previous study showed that Crif1 promotes adipogenic differentiation of BM-MSCs after radiation by modulating the cAMP/PKA signalling pathway(Zhang et al., 2015).

In this study, we investigated the role of Crif1 in osteoporosis after radiation. To address this question *in vivo*, we generated a *Crif1* bone marrow-specific conditional knockout mouse model. Conditional *Crif1* deletion causes decreases in RANKL expression and the RANKL/OPG ratio and reduces bone loss after radiation. In this study, we demonstrate that Crif1 promotes osteoporosis by regulating RANKL expression via the cAMP/PKA pathway. Through screening, we also identify five compounds that could effectively inhibit RANKL expression.

## Results

### Radiation induces osteoporosis in mice

To confirm the extent of bone loss over the short term after irradiation, we irradiated mice with a single dose of 5 Gy, and then, 7 days later, we harvested the left femurs. Micro-CT analysis of the distal femurs of males and females at 12 weeks of age revealed significant decreases in trabecular bone volume/total volume (BV/TV) (Figures 1A and 1B), connectivity density (Conn.D) (Figure 1C), trabecular number (Tb.N*) (Figure 1D), and bone mineral density (vBMD) (Figure 1E), but also showed significant increases in trabecular spacing (Tb.Sp*) (Figure 1G) and structure model index (SMI) (Figure 1H). There was no significant difference in trabecular thickness (Tb.Th*) (Figure 1F). Haematoxylin and eosin (H&E) staining of femoral sections from irradiated mice showed significantly decreased trabecular bone compared to controls (Figure 1I). Paraffin sections of femurs showed more TRAP-positive cells in irradiated mice than in control mice (Figure 1J). These results indicated that bone resorption was enhanced after radiation and validated the model of radiation-induced osteoporosis. Moreover, RT-qPCR data revealed dramatic increases in RANKL expression (Figure 1K) and the RANKL/OPG ratio in irradiated bone marrow cells (Figure 1L). OPG expression was not affected by radiation treatment (Figure 1K). Notably, expression of Crif1 also increased in irradiated bone marrow cells compared with control cells 7 days after irradiation (Figure 1M).

**Figure 1.**
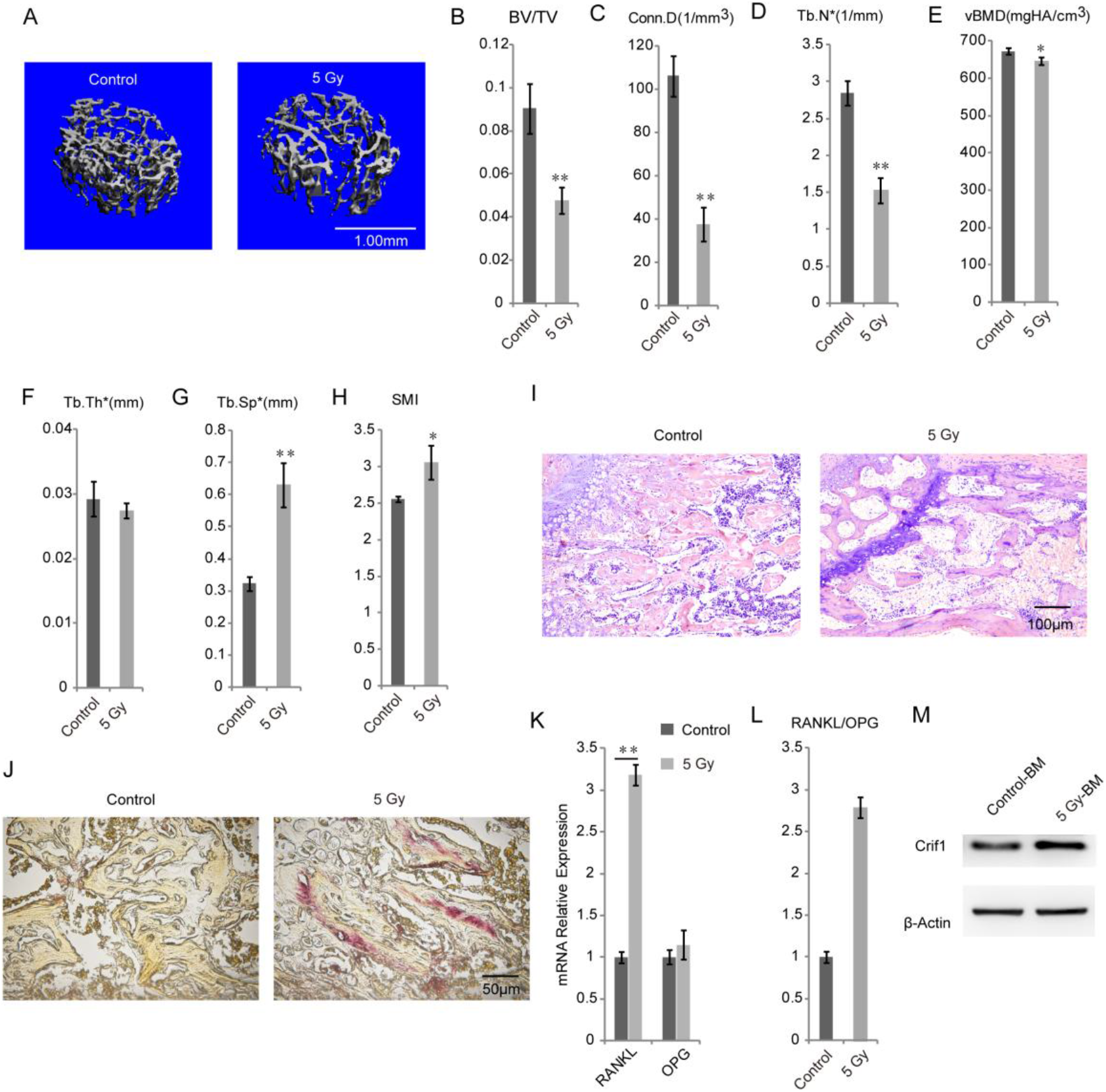
Radiation induces osteoporosis in mice. (A) Micro-CT images of the distal metaphysis of the femur. (B-H) Micro-CT analysis of the trabecular bone volume/total volume (B), connectivity density (C), trabecular number (D), bone mineral density (E), trabecular thickness (F), trabecular spacing (G), and structure model index (H). (I) H&E staining of femoral sections from irradiated mice and controls. (J) TRAP staining of femoral sections from irradiated mice and controls. RT-qPCR analysis of RANKL and OPG mRNA expression in flushed whole bone marrow. (K) RANKL/OPG ratio of RT-qPCR results. (L) Western blotting analysis of Crif1 expression in flushed whole bone marrow. *\*P < 0.05*, \*\**P* < 0.01. All experiments were performed in triplicate, and the bars represent the mean ± SD.

### Conditional Crif1 deletion from bone marrow relieves bone loss after radiation

To explore the possible relationship between Crif1 and RANKL, we constructed a bone marrow cell-specific Crif1-deficient mouse model by crossing *Crif*^*fl/fl*^ mice to *B6.129P2-Lyz2tm1(cre)/Nju* mice to generate *Lyz2Cre;Crif*^*fl/fl*^ mice (Figure 2A) and treated the mice with a single dose of 5 Gy. RT-qPCR results demonstrated that *Lyz2Cre;Crif*^*fl/fl*^ mice had lower RANKL expression (Figure 2B) and RANKL/OPG ratio in bone marrow cells after radiation compared to *Crif*^*fl/fl*^ mice (Figure 2C). OPG expression in bone marrow cells was not affected by *Crif1* deletion (Figure 2B). Therefore, we speculated that Crif1 may be involved in the regulation of RANKL expression. To further assess the effects of *Crif1* deletion on osteoporosis, trabecular bone parameters of femurs were evaluated by micro-CT analysis. Conditional *Crif1* deletion from bone marrow had no effect on bone mass (Figures 2D-2K). Although trabecular bone volume/total volume, connectivity density and trabecular number all decreased in *Crif*^*fl/fl*^ mice and *Lyz2Cre;Crif*^*fl/fl*^ mice after radiation compared to nonirradiated mice, these three parameters were significantly higher in irradiated *Lyz2Cre;Crif*^*fl/fl*^ mice than in irradiated *Crif*^*fl/fl*^ mice (Figures 2E,2F and 2G). Notably, trabecular spacing in *Crif*^*fl/fl*^ mice and *Lyz2Cre;Crif*^*fl/fl*^ mice both increased visibly after radiation, but the increase in *Lyz2Cre;Crif*^*fl/fl*^ mice was less than that of *Crif*^*fl/fl*^ mice after radiation treatment (Figure 2J). Furthermore, bone mineral density decreased (Figure 2H) and structural model index increased (Figure 2K) only in irradiated *Crif*^*fl/fl*^ mice compared to nonirradiated controls, while no remarkable changes in these parameters were detected between nonirradiated and irradiated *Lyz2Cre;Crif*^*fl/fl*^ mice (Figure 2H and 2K). In addition, trabecular thickness was not influenced by *Crif1* deletion or radiation treatment (Figure 2I). H&E staining of paraffin sections of femurs confirmed the previously reported micro-CT data depicting the reduced loss of trabecular tissue in *Lyz2Cre;Crif*^*fl/fl*^ mice compared to *Crif*^*fl/fl*^ mice after radiation (Figure 2L). We also detected a decreased number of TRAP-positive osteoclasts in irradiated *Lyz2Cre;Crif*^*fl/fl*^ mice compared to irradiated *Crif*^*fl/fl*^ mice, indicating weakened bone resorption (Figure 2M). Taken together, these data suggested that conditional *Crif1* deletion could alleviate bone resorption and reduce bone loss after radiation.

**Figure 2.**
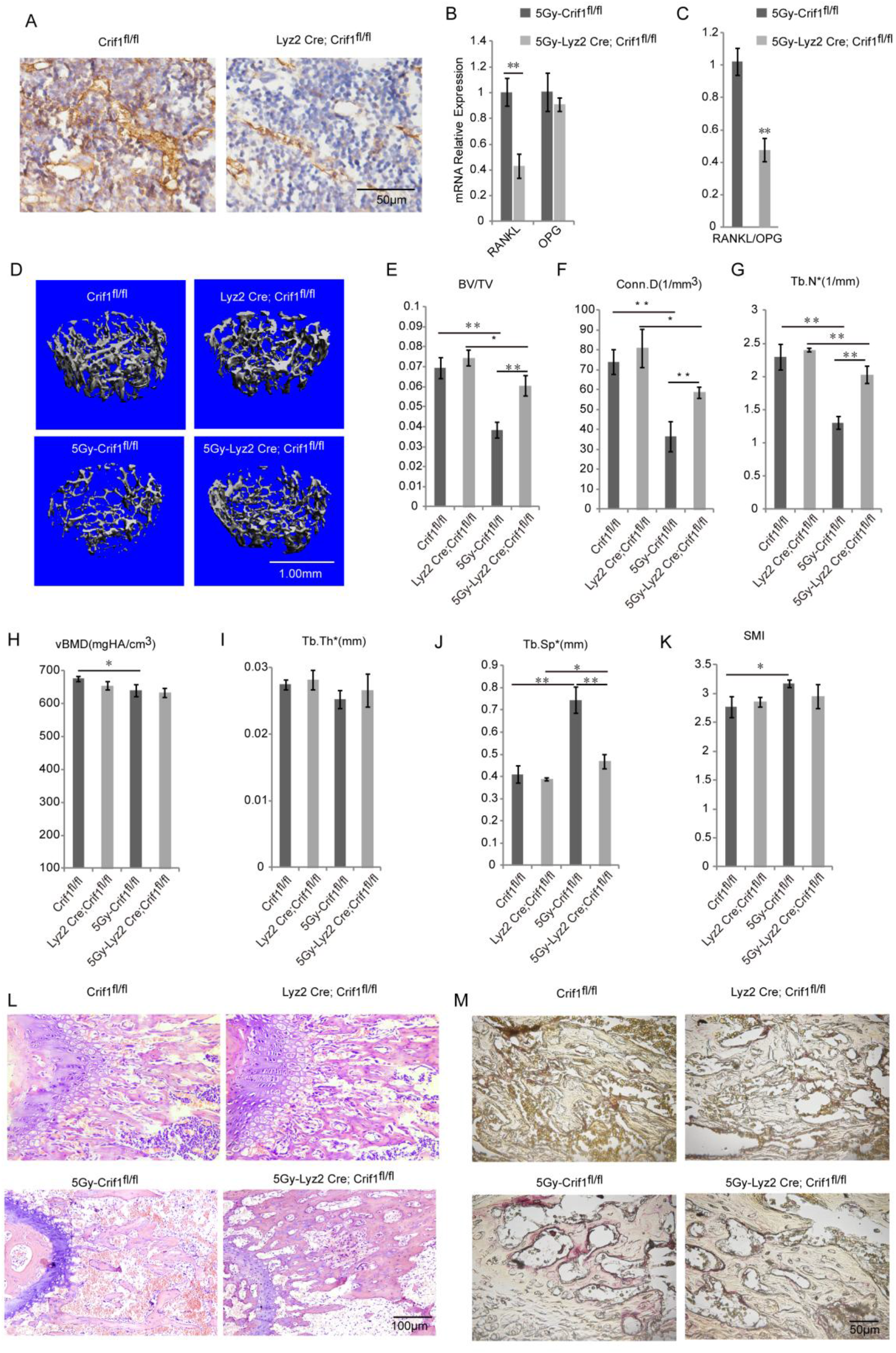
Conditional*Crif1* deletion from bone marrow reduces bone loss after radiation. (A) Histochemical staining analysis of Crif1 expression in femoral bone marrow. (B) RT-qPCR analysis of RANKL and OPG mRNA expression in flushed whole bone marrow of *Crif1*^*fl/fl*^ mice and *Lyz2Cre;Crif1*^*fl/fl*^ mice after 5 Gy radiation. (C) RANKL/OPG ratio of RT-qPCR results. (D)Micro-CT images of the distal metaphysis of the femur from *Lyz2Cre;Crif1*^*fl/fl*^ mice and *Crif1*^*fl/fl*^ mice. (E-K) Micro-CT analysis of the trabecular bone volume/total volume (E), connectivity density (F), trabecular number (G), bone mineral density (H), trabecular thickness (I), trabecular spacing (J), and structure model index (K). (L) H&E staining of femoral sections from *Crif1*^*fl/fl*^ mice and *Lyz2Cre;Crif1*^*fl/fl*^ mice. (M) TRAP staining of femoral sections from *Crif1*^*fl/fl*^ mice and *Lyz2Cre;Crif1*^*fl/fl*^ mice. \**P* < 0.05, \*\**P* < 0.01. All experiments were performed in triplicate, and the bars represent the mean ± SD.

### Overexpression of Crif1 in BM-MSCs increases RANKL secretion and the ratio of RANKL to OPG

To investigate the role of Crif1 in osteoporosis *in vitro*, we transfected mouse BM-MSCs with a Crif1 lentiviral overexpression vector (Figure 3A). For osteoclast induction *in vitro*, Crif1-overexpressing BM-MSCs and RAW264.7 cells were cocultured in a 12-well transwell unit for 7 days. RT-qPCR results showed that the relative mRNA expression of RANKL and RANKL/OPG ratio both increased in Crif1-overexpressing BM-MSCs compared to controls after 7 days of coculture (Figure 3B and 3C). Concentrations of RANKL and OPG in coculture medium were also detected by ELISA. Crif1-overexpressing BM-MSCs produced high levels of RANKL compared to the control (Figure 3D), while there was no significant difference in OPG concentration between the two groups (Figure 3E). The RANKL/OPG ratio in Crif1-overexpressing BM-MSCs was higher than in the control (Figure 3F). We also detected an increased number of tartrate-resistant alkaline phosphatase (TRAP)-positive cells in RAW264.7 cells cocultured with Crif1-overexpressing BM-MSCs (Figures 3G and 3H). These data suggested that Crif1 could promote RANKL expression and may be involved in osteoclast differentiation.

**Figure 3.**
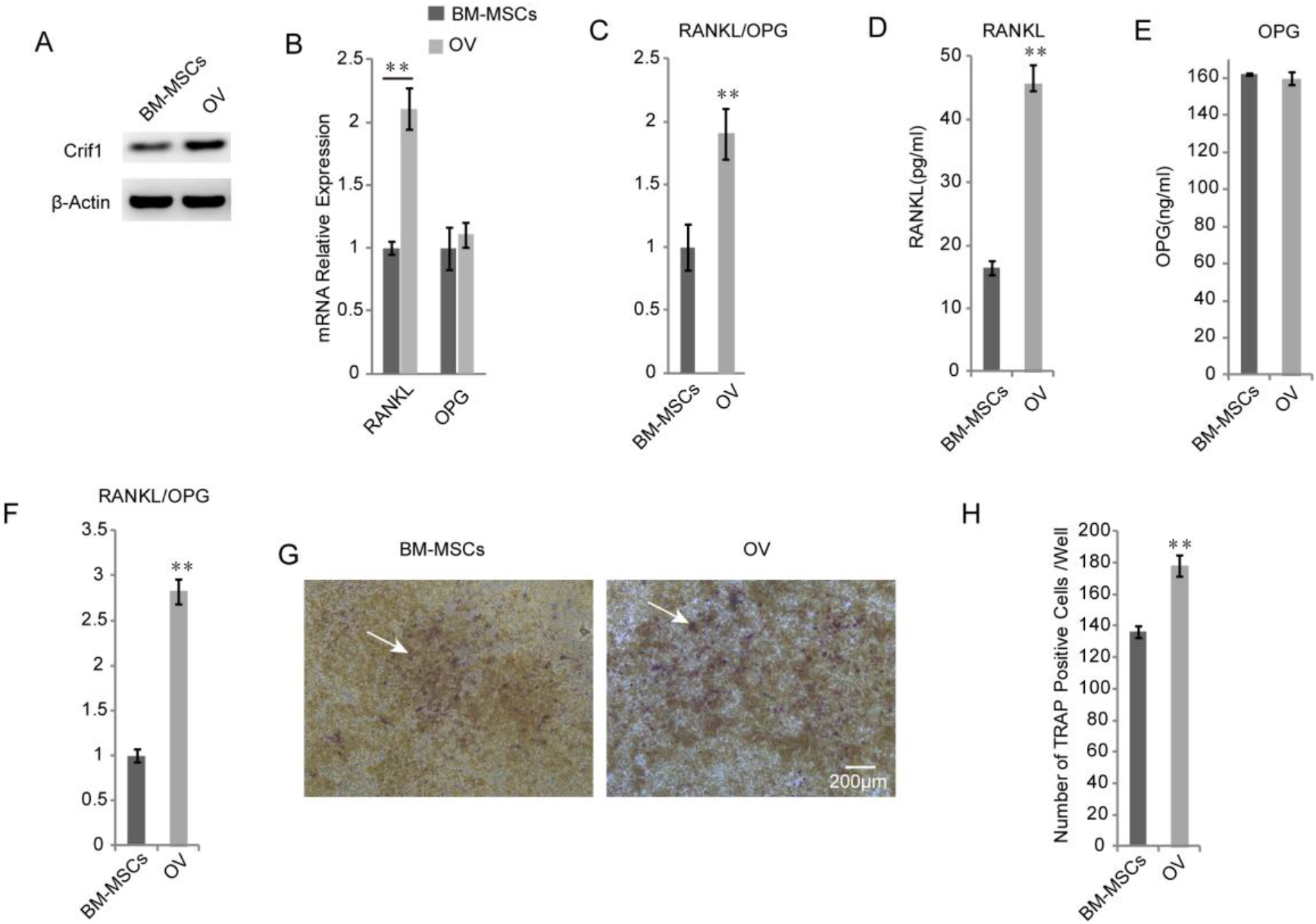
Overexpression of Crif1 in BM-MSCs results in increases in RANKL secretion and the ratio of RANKL to OPG. (A) Western blotting analysis of Crif1 expression in mouse BM-MSCs. Mouse BM-MSCs were transfected with a Crif1 lentiviral overexpression vector. (B) RT-qPCR analysis of RANKL and OPG mRNA expression in BM-MSCs and Crif1-overexpressing BM-MSCs (OV). BM-MSCs and OV were cocultured with RAW264.7. (C) RANKL/OPG ratio of RT-qPCR results. (D) ELISA analysis of RANKL protein levels in coculture supernatant medium. (E) ELISA analysis of OPG protein levels in coculture supernatant medium. (F)RANKL/OPG ratio in coculture supernatant medium. (G) TRAP staining of RAW264.7 after 7 days in coculture. (H) Average number of TRAP-positive cells/well (arrow) from RAW264.7 cells in coculture. *\*P < 0.05*, \*\**P* < 0.01. All experiments were performed in triplicate, and the bars represent the mean ± SD.

### Crfi1 is involved in the regulation of RANKL expression after radiation

To further confirm whether Crif1 played an important role in osteoporosis after radiation, we knocked out *Crif1* in RAW264.7 cells and BM-MSCs (Figures 4A and 4D), respectively. Deletion of *Crif1* from RAW264.7 cells did not affect RANK expression or osteoclast differentiation (Figures 4A, 4B and 4C). We previously demonstrated that Crif1 expression was upregulated after radiation(Zhang et al., 2015), and in this study, we found that RANKL expression and the RANKL/OPG ratio were also elevated after radiation (Figures 4E, 4F, 4G and 4I). Meanwhile, more TRAP-positive cells were found in RAW264.7 cells cocultured with BM-MSCs after radiation (Figures 4J and 4K). However, knocking out *Crif1* in BM-MSCs could significantly reduce RANKL expression and the RANKL/OPG ratio both before and after radiation (Figures 4E, 4F, 4G and 4I). OPG expression was not affected by *Crif1* deletion or radiation treatment (Figures 4E and 4H). Moreover, the number of TRAP-positive cells also decreased in *Crif1* knockout BM-MSCs compared to the control after 7 days of coculture (Figures 4J and 4K). These results further demonstrated that Crif1 can regulate RANKL expression, especially after radiation.

**Figure 4.**
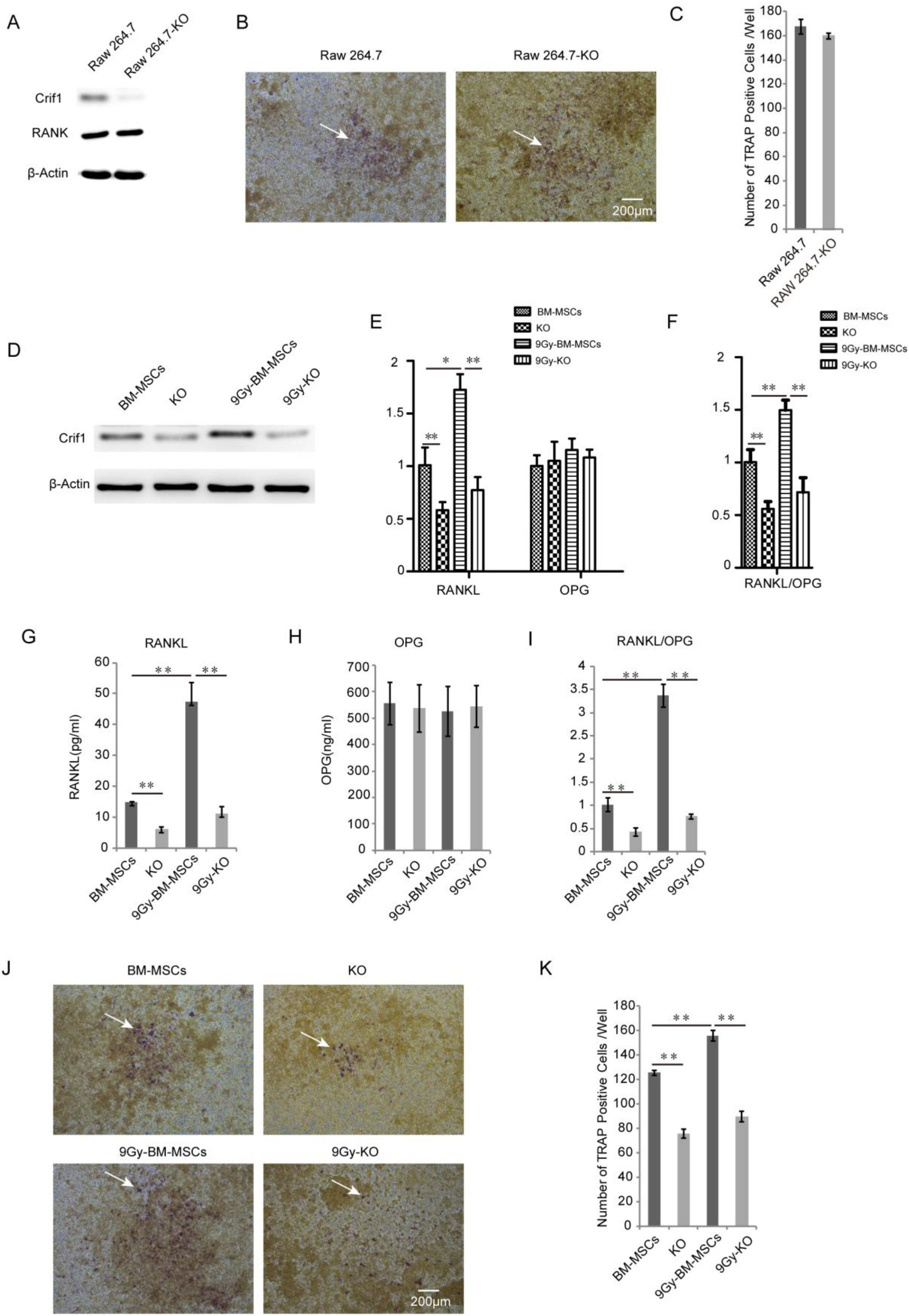
Crfi1 is involved in the regulation of RANKL expression after radiation. (A) Western blotting analysis of Crif1 and RANK expression in RAW264.7 cells. Crif1 was knocked out from RAW264.7 cells (RAW264.7-KO). (B) TRAP staining of RAW264.7-KO and controls after 7 days in coculture with mouse BM-MSCs. (C) Average number of TRAP-positive cells/well (arrow) from RAW264.7-KO and controls after 7 days in coculture with mouse BM-MSCs. (D) Western blotting analysis of Crif1 expression in BM-MSCs. Crif1 was knocked out from mouse BM-MSCs (KO), and KO and controls were irradiated with 9 Gy Co-60. (E) RT-qPCR analysis of RANKL and OPG mRNA expression in BM-MSCs and Crif1 knockout BM-MSCs (KO). BM-MSCs and KO were cocultured with RAW264.7 (F) RANKL/OPG ratio of RT-qPCR results. (G) ELISA analysis of RANKL protein levels in coculture supernatant medium. (H) ELISA analysis of OPG protein levels in coculture supernatant medium. (I) RANKL/OPG ratio in coculture supernatant medium. (J) TRAP staining of RAW264.7 after 7 days in coculture. (K) Average number of TRAP-positive cells/well (arrow) from RAW264.7 in coculture. *\*P < 0.05*, \*\**P* < 0.01. All experiments were performed in triplicate, and the bars represent the mean ± SD.

### Crif1 mediates adipogenesis and RANKL secretion in adipocytes

To determine whether Crif1 affects RANKL expression in adipocytes, BM-MSCs were grown in mouse mesenchymal stem cell adipogenic differentiation medium. Consistent with our previous research(Zhang et al., 2015), more BM-MCSs became strongly predisposed to adipogenesis (Figures 5A, 5B and 5C). Recently, it was reported that bone marrow adipocytes can secrete RANKL(Fan et al., 2017). Here, we found an obvious increase in RANKL expression and RANKL/OPG ratio in irradiated BM-MSCs after adipogenic induction (Figures 5D, 5E, 5F and 5H). However, knocking out *Crif1* in BM-MSCs reduced both adipogenesis and RANKL expression (Figures 5C, 5D and 5F). OPG expression was not affected by *Crif1* deletion or radiation treatment (Figures 5D and 5G). These data suggested that Crif1 mediates adipogenesis and RANKL secretion in adipocytes.

**Figure 5.**
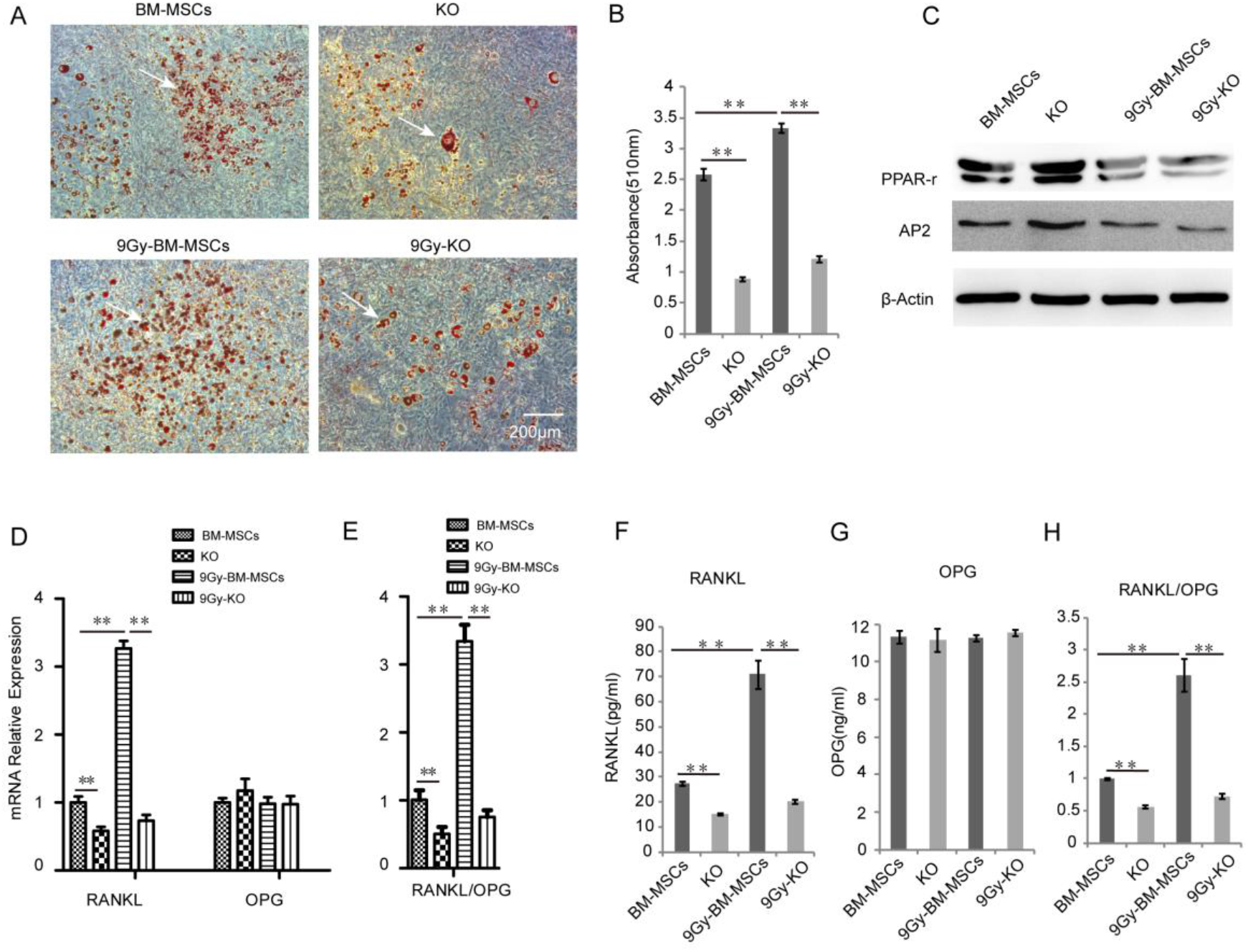
Crif1 mediates adipogenesis and RANKL secretion in adipocytes. (A) Oil red O staining analysis of mouse BM-MSCs after 21 days of adipogenic differentiation. Crif1 was knocked out from mouse BM-MSCs (KO), and KO and controls were irradiated with 9 Gy Co-60. KO and controls were treated with mouse mesenchymal stem cell adipogenic differentiation medium (Ad) to induce adipogenesis. (B) The dye from oil red O staining was extracted using isopropanol, and the optical density at 510 nm was measured using a Benchmark Plus. (C) Western blotting analysis of adipogenesis-related markers and transcription factors PPARγ and AP2 in mouse BM-MSCs after 21 days of adipogenic differentiation. (D) RT-qPCR analysis of RANKL and OPG mRNA expression in BM-MSCs and Crif1 knockout BM-MSCs (KO). (E) RANKL/OPG ratio of RT-qPCR results. (F) ELISA analysis of RANKL protein levels in supernatant adipogenic differentiation medium. (G) ELISA analysis of OPG protein levels in supernatant adipogenic differentiation medium. (H) RANKL/OPG ratio in supernatant adipogenic differentiation medium. *\*P < 0.05*, \*\**P* < 0.01..All experiments were performed in triplicate, and the bars represent the mean ± SD.

### Crif1 promotes RANKL secretion by modulating the cAMP/PKA signalling pathway

To verify the mechanism underlying Crif1-mediated upregulation of RANKL expression, PKA agonist (forskolin) and inhibitor (H89) were added to the coculture system. Although RANKL expression and the RANKL/OPG ratio were both increased after treatment with 25 μM forskolin, these effects were significantly weakened in Crif1 knockout BM-MSCs (Figures 6A, 6B, 6C and 6E). In addition, RANKL expression and the RANKL/OPG ratio were both decreased when Crif1-overexpressing BM-MSCs and controls were treated with 20 μM H89 (Figures 6F, 6G, 6H and 6J). OPG expression was not affected by forskolin or H89 treatment (Figures 6A, 6D, 6F and 6H). The most TRAP-positive cells were found in the coculture with forskolin-treated BM-MSCs (Figures 6K and 6L), the fewest TRAP-positive cells were found in the coculture with H89-treated cells (Figures 6M and 6N). After the addition of forskolin, CREB phosphorylation was significantly increased in the control BM-MSCs, but was dramatically inhibited in *Crif1* knockout BM-MCSs (Figure 6O). We also observed that CREB phosphorylation was suppressed in both Crif1-overexpressing BM-MSCs and controls following exposure to H89 (Figure 6P). These results demonstrated that Crif1 promotes RANKL expression through cAMP/PKA signalling pathway.

**Figure 6.**
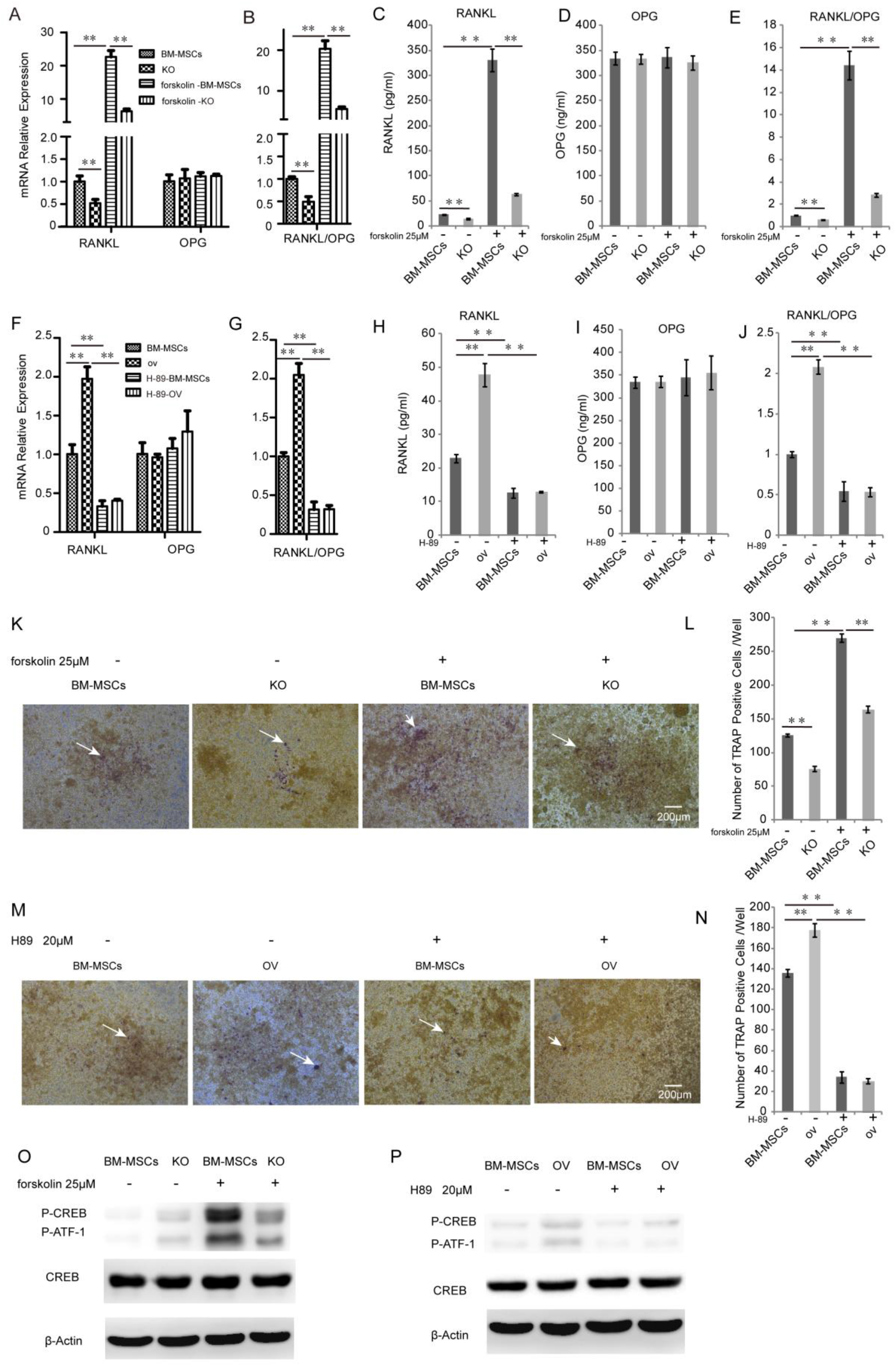
Crif1 promotes RANKL secretion by modulating the cAMP/PKA signalling pathway. (A)RT-qPCR analysis of RANKL and OPG mRNA expression in BM-MSCs and Crif1 knockout BM-MSCs (KO). BM-MSCs and KO were cocultured with RAW264.7 with or without 25 μM forskolin respectively. (B) RANKL/OPG ratio of RT-qPCR results. (C) ELISA analysis of RANKL protein levels in coculture supernatant medium with or without 25 μM forskolin. (D) ELISA analysis of OPG protein levels in coculture supernatant medium with or without 25 μM forskolin. (E) RANKL/OPG ratio in coculture supernatant medium with or without 25 μM forskolin. (F) RT-qPCR analysis of RANKL and OPG mRNA expression in BM-MSCs and Crif1-overexpressing BM-MSCs (OV). BM-MSCs and OV were cocultured with RAW264.7 with or without 20 μM H89 respectively. (G) RANKL/OPG ratio of RT-qPCR results. (H) ELISA analysis of RANKL protein levels in coculture supernatant medium with or without 20 μM H89. (I) ELISA analysis of OPG protein levels in coculture supernatant medium with or without 20 μM H89. (J) RANKL/OPG ratio in coculture supernatant medium with or without 20 μM H89. (K) TRAP staining of RAW264.7 in coculture treated with or without 25 μM forskolin. (L) Average number of TRAP-positive cells/well (arrow) from RAW264.7 in coculture treated with or without 25 μM forskolin. (M) TRAP staining of RAW264.7 in coculture treated with or without 20 μM H89. (N) Average number of TRAP-positive cells/well (arrow) from RAW264.7 in coculture treated with or without 20 μM H89. (O)Western blotting analysis of CREB phosphorylation levels in BM-MSCs in coculture treated with or without 25 μM forskolin. (P) Western blotting analysis of CREB phosphorylation levels in BM-MSCs in coculture treated with or without 20 μM H89. *\*P < 0.05*, \*\**P* < 0.01..All experiments were performed in triplicate, and the bars represent the mean ± SD.

### Crif1 inhibitors effectively suppress RANKL secretion and CREB phosphorylation

We used ClusPro and InterProSurf to investigate the interface in Crif1-PKAα complex, and results showed that Thr^197^, Gly^200^, Thr^201^, Glu^203^, and Phe^129^ of PKAα interact with Ile^132^, Met^128^, Ile^121^, His^120^, and Arg^117^ in the long alpha helical region of Crif1, forming the binding interface (Figure 7A). A virtual screening using 462,608 compounds from the Life Chemicals database around His^120^ of Crif1 was carried out using the program Autodock_vina. A set of 13 compounds was selected for experimental screening based on binding energy <-12.0 kcal/mol (Table S1). Initially a tetrazolium salt (WST-8) assay was carried out to study potential toxic effects of these compounds on hBM-MSCs, vero cells and mouse BM-MSCs (Figures S1-S3). The compounds F0382-0033, F3408-0076, F1430-0134, F3408-0031 and F1430-0130 showed low toxicity to hBM-MSCs at concentrations of 25 μM (Figure 7G-7K). The binding pattern in these ligand−protein complexes potentially contained multiple interactions dominated by hydrophobic amino acids (Figure 7B-7F). To determine whether these 5 Crif1 inhibitors affected RANKL expression, hBM-MSCs were pretreated with these compounds followed by treatment with forskolin. ELISA analysis of supernatant medium revealed that RANKL expression was dramatically decreased by treatment with Crif1 inhibitors (Figure 7L). OPG expression could be significantly increased by F1430-0134 (Figure 7M). Moreover, RANKL/OPG ratios were also decreased by these 5 inhibitors (Figure 7N). To further understand the mechanism, CREB phosphorylation was detected. Western blotting analysis showed that CREB phosphorylation was inhibited by treatment with the 5 inhibitors that showed suppressive effects on RANKL expression (Figure 7O).

**Figure 7.**
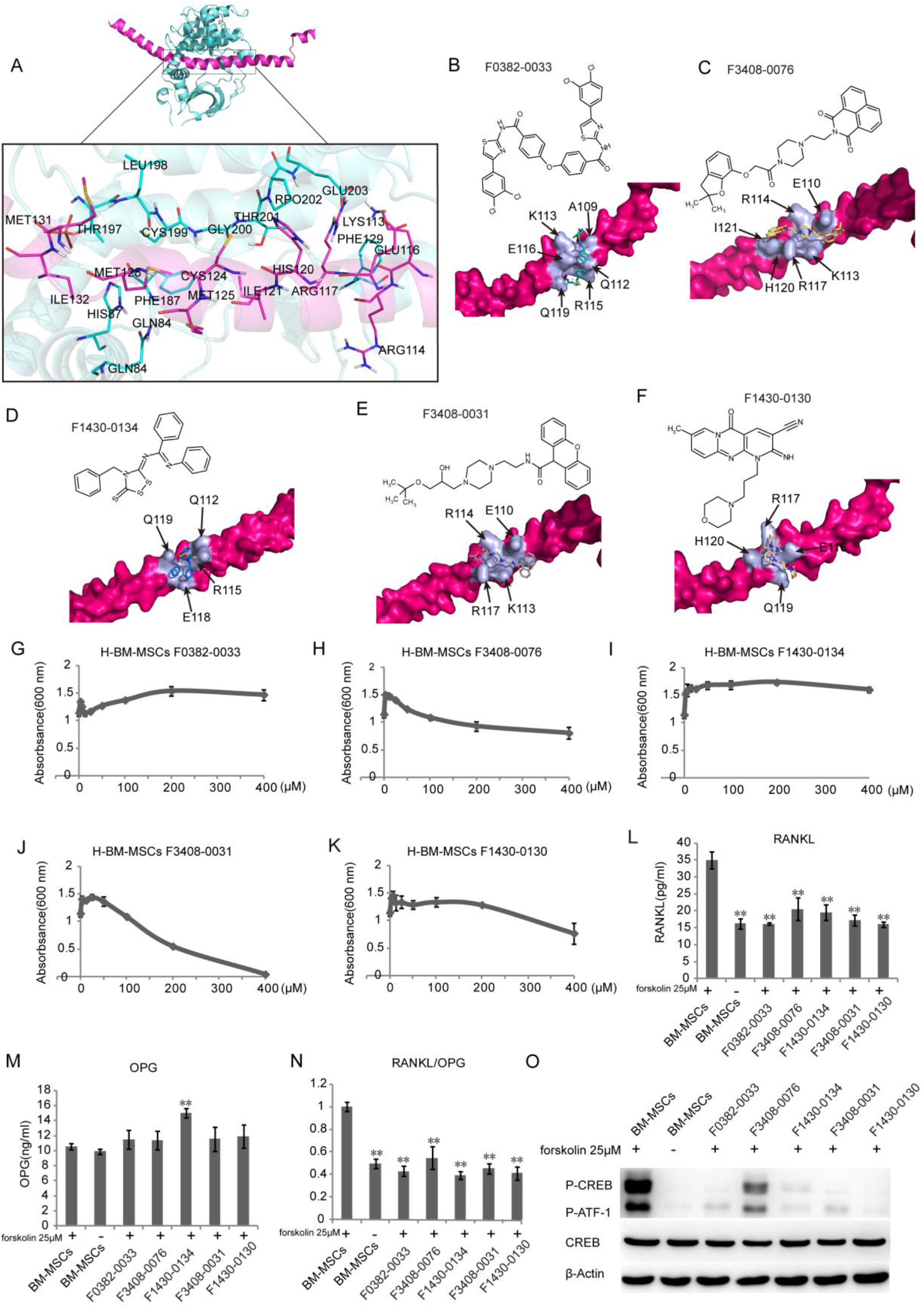
Crif1 inhibitors effectively suppress RANKL secretion and CREB phosphorylation. (A) Crif1− PKAα interaction model showing Crif1 (colored in rose red) and PKAα (colored in cyan). Interface amino acids are shown as sticks and colored in rose red (for Crif1) and cyan (for PKAα) and indicated as a zoomed-in view in the inset figure. (B-F) Chemical structure of each inhibitor molecule and their docked pose on Crif1 (colored in rose red, surface view). Docked molecule (stick) and the amino acids involved in the hydrophobic interactions (light purple) are shown. F0382-0033 (B), F3408-0076 (C), F1430-0134 (D), F3408-0031 (E) and F1430-0130 (F). (G-K) A tetrazolium salt (WST-8) assay was carried out to study the toxity effect of compounds on the hBM-MSCs. F0382-0033 (G), F3408-0076 (H), F1430-0134 (I), F3408-0031 (J) and F1430-0130 (K). (L)ELISA analysis of RANKL protein levels in supernatant medium. hBM-MSCs were pretreated with 5 different compounds followed by treatment with forskolin, and supernatant medium was collected for ELISA after 3 days. (M) ELISA analysis of OPG protein levels in supernatant medium. (N) RANKL/OPG ratio in supernatant medium. (O)Western blotting analysis of CREB phosphorylation levels. hBM-MSCs were pretreated with 5 different compounds followed by treatment with forskolin and total protein lysates were extracted for CREB phosphorylation detection after 1 hour. *\*P < 0.05, **P < 0.01*. All experiments were performed in triplicate, and the bars represent the mean ± SD.

## Discussion

Radiation exposure (due to radiotherapy, accidental causes or terrorism) causes irreparable damage to tissues and organs such as bone marrow. It suppresses bone formation and elevates resorption, disturbing the balance between them and leading to a dramatic decline in trabecular bone(D’Oronzo, Stucci, Tucci, & Silvestris, 2015; Jia, Gaddy, Suva, & Corry, 2011). In this study, we treated mice with a single dose of 5 Gy to generate a model of radiation-induced osteoporosis. RANKL expression and RANKL/OPG ratios actually increased in the surviving bone marrow cells after radiation, which was consistent with a previous study(Alwood et al., 2015). Meanwhile, expression of Crif1 and bone resorption also increased, indicating a relationship between RANKL and Crif1 in osteoporosis. BM-MSCs, osteoblasts, adipocytes and activated T lymphocytes in bone marrow are the main source of RANKL secretion(Boyce & Xing, 2008; Honma et al., 2013); thus, to elucidate the possible role of Crif1 in RANKL expression, we generated bone marrow cell-specific Crif1-deficient mice. Conditional *Crif1* deletion from bone marrow could reduce RANKL expression and relieve bone loss after radiation, suggesting that Crif1 is involved in RANKL expression and osteoporosis.

BM-MSCs, which are a major and important component of the haematopoietic microenvironment, secrete soluble RANKL and OPG(Horowitz, Xi, Wilson, & Kacena, 2001; Morrison & Scadden, 2014). In our study, overexpression of Crif1 in BM-MSCs resulted in an increase in RANKL secretion, while deletion of Crif1 from BM-MSCs significantly reduced RANKL expression. BM-MSCs are pluripotent and are the progenitors of both bone marrow osteocytes and adipocytes. The balance between osteogenic and adipogenic differentiation of BM-MSCs also plays a pivotal role in maintaining bone homeostasis(Mori et al., 2014; Yue, Zhou, Shimada, Zhao, & Morrison, 2016). Excessive numbers of adipocytes are often found in the bone marrow of patients with osteoporosis. A recent study revealed that bone marrow adipocytes can also secrete RANKL and accelerate osteoclastogenesis(Fan et al., 2017). Moreover, after radiation exposure, the haematopoietic red marrow gradually turns to yellow, a phenomenon known as bone marrow fatting. Compared to other cells in bone marrow, BM-MSCs are more resistant to radiation; however, radiation impairs the balance between osteogenic and adipogenic differentiation of BM-MSCs, decreasing osteogenesis while increasing RANKL expression and adipogenesis(Cao et al., 2011). We previously reported that Crif1 can promote the adipogenesis of BM-MSCs. Here, we found that deletion of Crif1 from BM-MSCs can reduce both adipogenesis and RANKL expression. Osteogenesis and adipogenesis in the bone marrow are inversely correlated, so reduced adipogenesis results in an increase in the osteoblast pool(Muruganandan, Roman, & Sinal, 2009; Nuttall & Gimble, 2000).

RANKL functions as an osteoclast-activating factor, and its binding to RANK induces the activation of transcription factors such as c-fos, NFAT, and nuclear factor kappa B (NF-κB) in preosteoclasts and initiates several downstream signalling pathways, especially the NF-κB pathway(Nakashima, Hayashi, & Takayanagi, 2012). RANKL expression can be upregulated by many agents, such as PTH and forskolin. Forskolin can stimulate RANKL expression through the cAMP/PKA signalling pathway(Kondo, Guo, & Bringhurst, 2002; Tseng et al., 2010). However, deletion of Crif1 from BM-MSCs impairs the promotion of RANKL expression by forskolin. Moreover, overexpression of Crif1 in BM-MSCs does not increase RANKL expression upon exposure to a PKA inhibitor. Our previous study revealed that Crif1 can interact with PKA and promote adipogenesis. Here, we further demonstrated that Crif1 can also promote RANKL expression through the cAMP/PKA signalling pathway.

Drugs for the treatment of osteoporosis could be divided into anabolic and antiresorptive categories. Bisphosphonates (including alendronate and ibandronate), oestrogen, selective oestrogen receptor modulators (SERMs), and denosumab are antiresorptive drugs, whereas parathyroid hormone (PTH) and its analogues are anabolic agents(Andreopoulou & Bockman, 2015; Favus, 2010; Torstrick & Guldberg, 2014). However, the long-term adverse effects of these anti-osteoporotic drugs should be considered, such as hypocalcaemia, arthralgia, nausea, and especially the development of breast cancer and risks of cardiovascular events and thromboembolism associated with treatment with oestrogen and SERMs(Black & Rosen, 2016; Khan, Cheung, & Khan, 2017). Therefore, it is necessary to look for new drugs with high efficiency but few side effects. Because of the importance of RANKL in osteoclast differentiation, RANKL-secreting cells, which play a central role in osteoclastogenesis, are the targets of most antiresorptive agents(Dempster, Lambing, Kostenuik, & Grauer, 2012; Tella & Gallagher, 2014). Future drug screening should target not only the regulation of the balance between bone formation and bone resorption but also the balance between osteogenic and adipogenic differentiation. Here, through screening, we identified five compounds that effectively inhibit Crif1 activity in the cAMP/PKA signalling pathway. These five compounds may have dual actions on bone metabolism, both increasing bone formation and decreasing bone resorption and adipogenesis.

In summary, we demonstrated that Crif1 plays a crucial role in osteoporosis and that conditional *Crif1* deletion reduces bone loss after radiation. Moreover, through screening, we identified five Crif1 inhibitors that could dramatically reduce RANKL secretion and CREB phosphorylation. This study enriches current knowledge of the pathogenesis of radiation-induced osteoporosis and provides insights into potential therapeutic strategies for osteoporosis treatment.

## Materials and Methods

### Mouse Strains

Male and female C57BL/6 mice (aged 12-14 weeks) were purchased from Beijing HFK Bio-Technology Co. Ltd. *B6.129P2-Lyz2tm1(cre)/Nju* mice were received as a gift from Assistant Professor Tao Wang (Third Military Medical University). *Crif*^*fl/fl*^ mice were constructed by Nanjing BioMedical Research Institute of Nanjing University (NBRI). *Lyz2Cre;Crif1*^*fl/fl*^ mice were generated by crossing *Crif*^*fl/fl*^ mice to *B6.129P2-Lyz2tm1(cre)/Nju* transgenic mice. For radiation treatment, mice were irradiated with 5 Gy Co-60 at a rate of 0.69 Gy/minute. Blood was collected at 7 days after irradiation, and serum from each mouse was analysed individually in triplicate.

### Cell culture and treatment

For *in vitro* study, mouse bone marrow mesenchymal stem/stromal cells (BM-MSCs) purchased from Cyagen Biosciences were cultured in mouse mesenchymal stem cell medium (MUCMX-90011, Cyagen Biosciences) at 37°C and 5% CO_2_. For radiation treatment, mouse BM-MSCs were irradiated with 9 Gy Co-60 at a rate of 0.69 Gy/minute.

RAW264.7 cells were cultured in Dulbecco’s modified Eagle medium (DMEM, HyClone) supplemented with 10% FBS.

For osteoclast induction, RAW264.7 cells and mouse BM-MSCs were cocultured in a 12-well transwell unit (0.4 μm) for 7 days with or without forskolin (25 μM) or H89 (20 μM) treatment. After 7 days of coculture, cells were collected for RT-qPCR and Western blotting; meanwhile, supernatant medium was collected for ELISA.

Human bone marrow mesenchymal stem/stromal cells (hBM-MSCs) (catalogue No. 7500, ScienCell) were cultured in mesenchymal stem cell medium (catalogue No. 7501, ScienCell) at 37 °C and 5% CO_2_.

### *Crif1* knockout and overexpression in vitro

For Crif1 overexpression, mouse BM-MSCs were transfected with a Crif1 lentiviral overexpression vector (pLV[Exp]-EGFP:T2A:Puro-EF1A>mGadd45gip1 [NM_183358.4]) constructed by Cyagen Biosciences (vector ID: VB180112-1182ypt) and selected with 5 μg/ml puromycin dihydrochloride (A1113803, Invitrogen). An empty vector (pLV[Exp]-EGFP:T2A:Puro-Null, vector ID: VB160420-1011mqh, Cyagen Biosciences) was included as a control.

For *Crif1* knockout, mouse BM-MSCs were first transfected with lentiCas9-Blast vector (Genomeditech) and selected with 5 μg/ml blasticidin S HCl (A1113903, Invitrogen). Then, cells were transfected with CRISPR/Cas9 M_Gadd45gip1 gRNA vector (target sequence: GCGGGGGCGCACGGTAGCTG, Genomeditech) and selected with 5 μg/ml puromycin dihydrochloride. An empty vector (LentiGuide-Puro-Scramble-gRNA, Genomeditech) was included as a control.

### *In vitro* adipogenic differentiation

To induce adipogenesis, mouse BM-MSCs were treated in mouse mesenchymal stem cell adipogenic differentiation medium (MUCMX-90031, Cyagen Biosciences). After differentiation, we preserved the supernatant medium for ELISA and fixed the cells with 2 ml of 4% formaldehyde solution for 30 minutes. Then, the cells were stained with 1 ml oil red O working solution (catalogue No. S0131, Cyagen Biosciences) for 30 minutes and visualized with a light microscope (Leica DMIRB, Heidelberg, Germany). The dye from oil red O staining was extracted using isopropanol, and the optical density at 510 nm was measured using a Varioskan FLASH microplate reader.

### Micro-CT analysis

Femurs were dissected, fixed overnight in 4% paraformaldehyde, and stored in 1% paraformaldehyde at 4°C. Trabecular bone parameters were measured in the distal metaphysis of the femur. We started analysing slices at the bottom of the distal growth plate, where the epiphyseal cap structure completely disappeared and continued for 95 slices (10.5 μm/ slice, using SCANCO VivaCT40) towards the proximal end of the femur.

### Isolation of bone marrow cells

Femurs were collected and cleaned in sterile PBS, and both ends of each femur were trimmed off. Bones were placed in a 0.6-mL microcentrifuge tube that was cut open at the bottom and nested inside a 1.5-mL microcentrifuge tube. Fresh bone marrow was spun out by brief centrifugation (from 0 to 10,000 rpm, 9 s). Red blood cells were lysed using RBC lysis buffer (catalogue NO. RT122-02, TIANGEN). After centrifugation (3,000 rpm, 5 minutes), cells at the bottom layer were collected for Western blotting and RT-qPCR assays(Fan et al., 2017).

### Western blotting and antibodies

Protein expression in the samples was analysed by Western blotting. Total protein lysates were extracted with cell lysis buffer for Western blotting and immunoprecipitation (catalogue No. P0013, Beyotime) and denatured by boiling. Protein samples were resolved on 12% SDS-polyacrylamide gels and transferred to polyvinylidene fluoride membranes (PVDF Western Blotting Membranes; Roche). Membranes were blocked in PBS containing 5% (w/v) nonfat dry milk and 0.1% TWEEN 20 and then incubated with the appropriate primary antibodies overnight at 4°C. Membranes were washed with TBST three times, and then incubated with the appropriate horseradish peroxidase-conjugated secondary antibodies for 1 h at 24°C. Immunoreactive bands were detected by the BeyoECL Plus reagent (P0018, Beyotime) using a Photo-Image System (Molecular Dynamics, Sunnyvale, CA, USA). The primary antibodies used for blotting were as follows: Crif1 (M-222) (sc-134882; Santa Cruz), RANK (H-7) (sc-374360; Santa Cruz), A-FABP (AP2, sc-18661; Santa Cruz), PPARγ (sc-7273; Santa Cruz), β-actin (sc-47778; Santa Cruz), phospho-CREB rabbit mAb (#9198; Cell Signaling Technology), and CREB rabbit mAb (#9197; Cell Signaling Technology).

### RT-qPCR

Real-time quantitative polymerase chain reaction (RT-qPCR) was used to analyse the mRNA levels of selected genes in collected samples. Total RNA was extracted using TRIzol Reagent (catalogue No. 10296010, Invitrogen) according to the manufacturer’s instructions. First-strand cDNA was synthesized from l μg of RNA using the PrimeScript RT Reagent Kit with gDNA Eraser (catalogue No. RR047A, TaKaRa). qPCR was performed in triplicate in 20-μl reactions containing SYBR Premix Ex Taq II (catalogue No. RR820A, TaKaRa). The reaction protocol was as follows: heating for 30 s at 95°C, followed by 40 cycles of amplification (5 s at 95°C, 30 s at 60°C).

The sequences of the RT-PCR primers were as follows:

M-actin-F: AGCCATGTACGTAGCCATCC
M-actin-R: CTCAGCTGTGGTGGTGAA
M-Rankl-F: CTCCGAGCTGGTGAAGAAA
M-Rankl-:CCCCAAAGTACGTCGCATCT
M-OPG-F: GTTCCTGCACAGCTTCACAA
M-OPG-R: AACAGCCCAGTGACCATTC.

### ELISA

The concentrations of RANKL and OPG were measured using the Mouse RANκL (Receptor Activator of Nuclear Factor Kappa B Ligand) ELISA Kit (E-EL-M0644c, elabscience), Human sRANKL (Soluble Receptor Activator of Nuclear factor-kB Ligand) ELISA Kit (E-EL-H5558c, elabscience), Mouse OPG (Osteoprotegerin) ELISA Kit (E-EL-M0081c, elabscience) and Human OPG (Osteoprotegerin) ELISA Kit (E-EL-H1341c, elabscience) according to the manufacturer’s instructions. Assays were performed in triplicate.

### Tartrate-resistant acid phosphatase (TRAP) staining

After the 7-day coculture period, cells were washed once with PBS, fixed in 10% formalin for 10 minutes, and incubated with a substrate solution, naphthol AS-BI phosphate (catalogue No. 387, Sigma), in the presence of 50 mM sodium tartrate at 37 °C for 1 h. The resulting mononuclear and multinuclear TRAP-positive cells were visualized by light microscopy and quantified.

### Histochemistry and histomorphometric analysis

Femurs were dissected, fixed overnight in 4% paraformaldehyde, decalcified in 10% EDTA (pH 7.0) for 20 days and embedded in paraffin. Longitudinally oriented sections of bone 4 μm thick, including the metaphysis and diaphysis, were processed for histochemistry and haematoxylin and eosin (H&E) staining. Dewaxed sections were also stained for tartrate-resistant acid phosphatase (TRAP) activity to identify osteoclasts. Sections were incubated in TRAP stain for 45 minutes at 37°C.

### Crif1 inhibitor screening

ClusPro and InterProSurf were used to investigate the interface in Crif1-PKAα complex. A virtual screening using 462,608 compounds from the Life Chemicals database around His^120^ of Crif1 was carried out using the program Autodock_vina. For inhibitor screening, hBM-MSCs were cultured in 6-well plates and pretreated with 5 different compounds (25 μM). After 3 hours of pretreatment, forskolin (25 μM) was added to the medium. After 1 hour of forskolin treatment, total protein lysates were extracted for CREB phosphorylation detection, and 3 days later, supernatant medium was collected for ELISA.

### Tetrazolium salt (WST-8) assay

A tetrazolium salt (WST-8) assay was carried out to study the toxicity of compounds to hBM-MSCs. hBM-MSCs seeded at a density of 3000 cells per well in 96-well plate were treated with 5 different compounds at 8 final concentrations (3.125μM, 6.25μM, 12.5μM, 25μM, 50μM, 100μM, 200μM, 400μM). Three days later, 10μl cell counting kit -8 solution was added to each well. After 4 hours of incubation, the absorbance at 450 nm was measured using a Varioskan FLASH microplate reader (Thermo).

### Statistical analysis

The mRNA expression levels of RANKL and OPG in the tested samples were determined as the cycle threshold (CT) level, and normalized copy numbers (relative quantification) were calculated using the ΔΔCT equation as follows: ΔΔCT=ΔCT of the bone marrow sample -ΔCT of β-actin, and then the normalized copy number (relative quantification)=2^−^ΔΔ^CT^. Data are presented as the mean±SD. Statistical significance was assessed using a two-tailed paired Student’s t-test. The results were considered significant when * p<0.05 or ** p<0.01.

### Study approval

All animal studies performed were approved by the Institutional Animal Care and Use Committee at the Third Military Medical University.

## Acknowledgements

This work was supported by the National Natural Science Foundation of China (no.81502754), the Interdisciplinary and International Cooperation Projects of The Second Affiliated Hospital, Third Military Medical University (no. 2016YXKJC0) and the National Natural Science Foundation of China (no.31571352).

## Author Contributions

Z.L., L.X., Q.R. and L.C.: conceived the project; L.X. and Q.R.: designed and performed most experiments and data analysis with input; Y.X., F.L., M.Z., L.C., Y.X., L.Z., J.F.Z. and S.C.L.: assisted with experiments; L.X.: wrote and edited the manuscript.

## Supplementary data

**Table S1.** 13 compounds were chosen based on binding energy <−12.0 kcal/mol.

| Compound Name | Affinity (kcal/mol) |
| --- | --- |
| **F3408-0076** | -14.6 |
| **F0382-0033** | -13.1 |
| **F1430-0130** | -12.9 |
| **F0808-0153** | -12.5 |
| **F0808-0131** | -12.5 |
| **F0808-0151** | -12.4 |
| **F0808-1181** | -12.4 |
| **F0808-1175** | -12.4 |
| **F1269-0120** | -12.3 |
| **F0808-0152** | -12.2 |
| **F3408-0031** | -12.2 |
| **F1430-0134** | -12 |
| **F1430-0138** | -12 |

**Figure S1.**
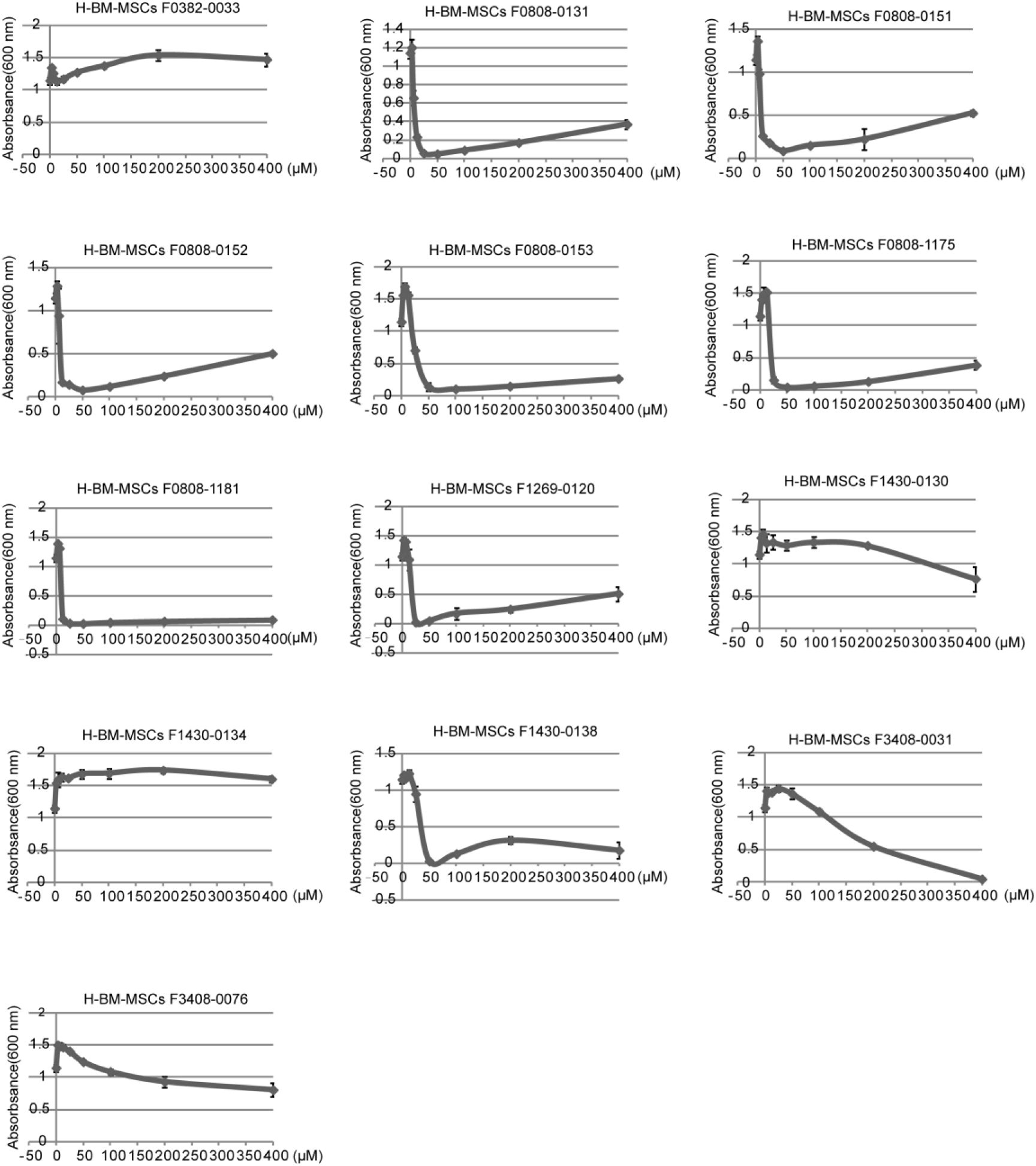
A tetrazolium salt (WST-8) assay was carried out to study the toxic effects of compounds on H-BM-MSCs.

**Figure S2.**
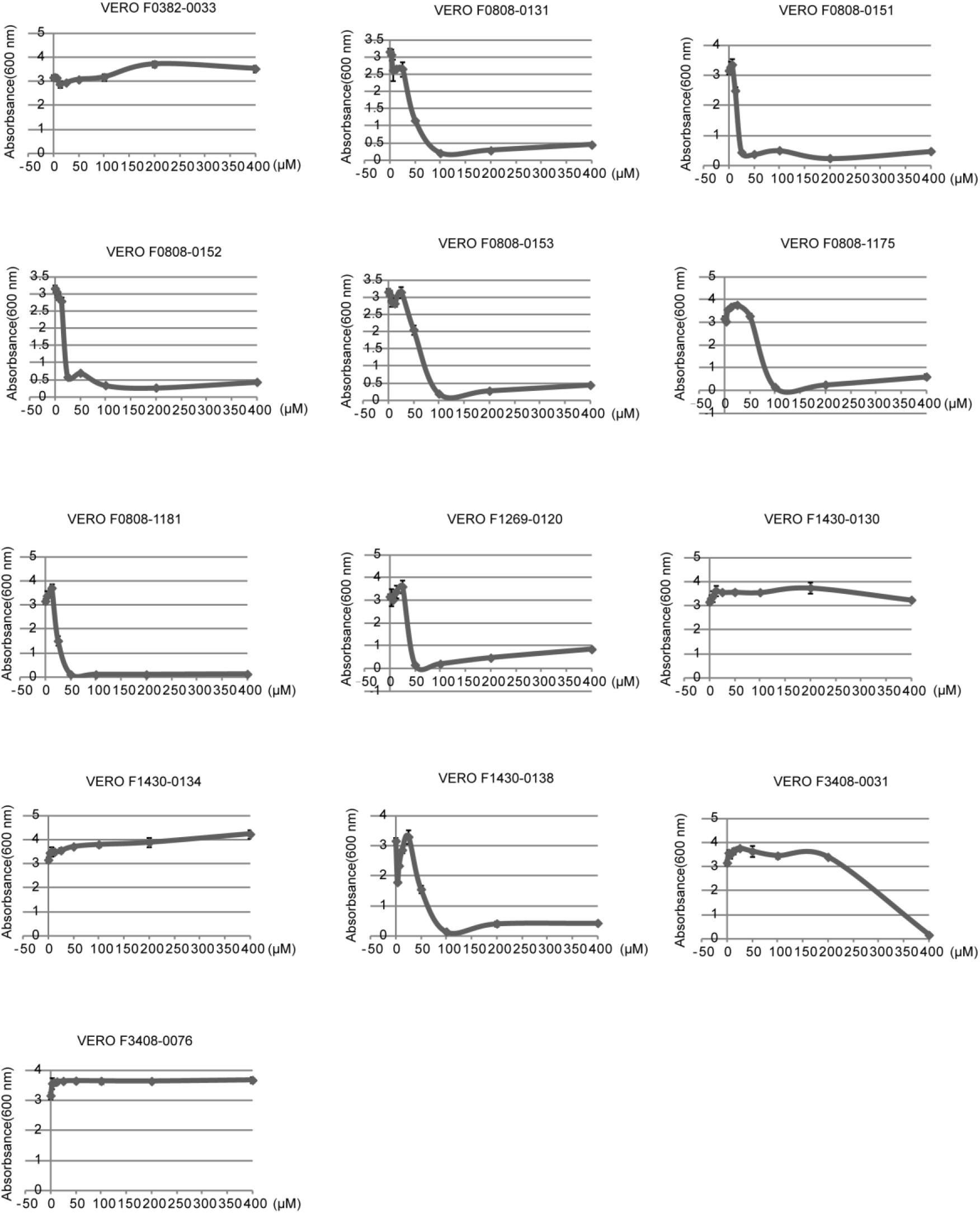
A tetrazolium salt (WST-8) assay was carried out to study the toxic effects of compounds on vero cells.

**Figure S3.**
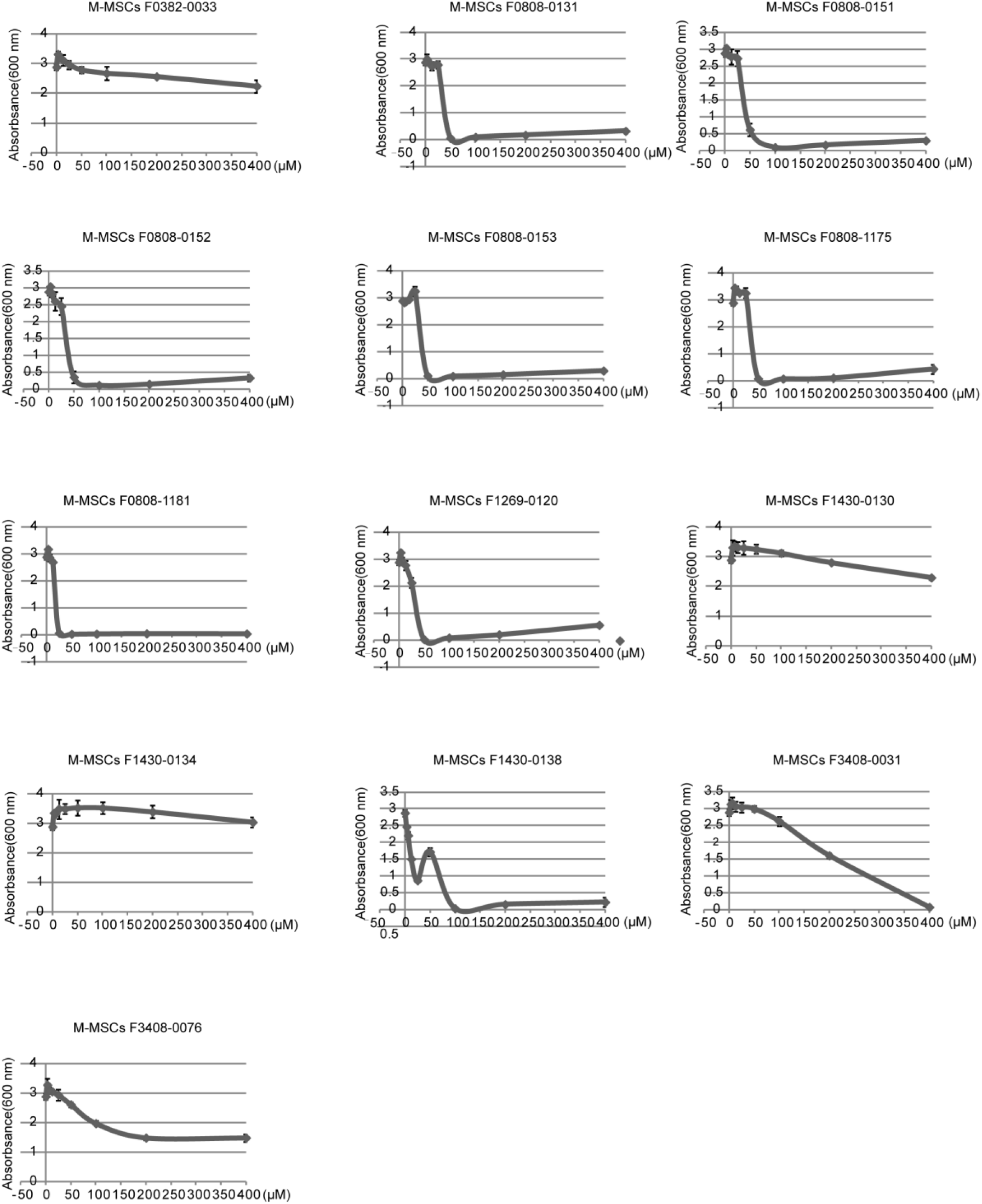
A tetrazolium salt (WST-8) assay was carried out to study the toxic effects of compounds on mouse BM-MSCs.

